# Ultrafast fMRI detects age-related changes in harmonics of cardiac pulsations in the brain at 7 T

**DOI:** 10.1101/2025.10.28.685077

**Authors:** Charles Marchini, Dema Abdelkarim, Yuhui Chai, Monica Fabiani, Gabriele Gratton, Bradley P. Sutton

**Affiliations:** Beckman Institute for Advanced Science and Technology, University of Illinois at Urbana-Champaign; Grainger College of Engineering, Bioengineering Department, University of Illinois at Urbana-Champaign; Carle Illinois Advanced Imaging Center, Carle Hospital and University of Illinois at Urbana-Champaign; Neuroscience Program, University of Illinois at Urbana-Champaign, Champaign, IL, USA; Department of Psychology, University of Illinois at Urbana-Champaign, Champaign, IL, USA; Carle Illinois College of Medicine, University of Illinois at Urbana-Champaign

**Keywords:** cerebrovascular pulsatility, partial separability, 7 Tesla, BOLD fMRI, aging, arterial stiffening

## Abstract

**Purpose:** To develop and apply an ultra-fast fMRI acquisition at 7 T to measure the pulsatile signals in the brain with full brain coverage and detect age-related differences. Pulsatile signals’ parameters provide information about the health status of the cerebrovascular system. This new acquisition provides a good mix of spatial coverage, image resolution, and temporal resolution for observing physiological signals.

**Methods:** A partial separability MRI acquisition and reconstruction approach was used to collect 3D data acquired at 7 T with TR = 65 ms in 28 healthy adults. 8 young subjects and 14 older subjects (10 male and 12 female) had acceptable pulse plethysmography and MRI data. With full-brain coverage and 2 mm isotropic resolution, the reliability of the pulse signal (even/odd heartbeat as test/retest) within each brain voxel was computed across subjects. Within voxels of high pulse reliability, the average magnitude within 0.15 Hz of the 4 heartbeat harmonic frequencies were compared within CSF, GM, and WM between young/old and between male/female participants.

**Results:** Increased first harmonic magnitude in older adults compared to young adults was detected in regions with reliable cardiac pulsations, which overlap major arteries and CSF pools. There was also a greater magnitude of the third harmonic normalized to the first in males. Brain maps of the pulse frequency and the harmonics were formed for visualization of regions of high pulsatility.

**Conclusion:** This new approach captures the pulsatile signal with BOLD fMRI with high spatiotemporal resolution and can be sensitive to age- and sex-related differences.

## 1 INTRODUCTION

Aging is associated with changes in cognitive and brain function, which can lead to the development of mild cognitive impairment and ultimately dementia. Risk factors for these outcomes largely overlap those for cardiovascular and cerebrovascular disease. Epidemiologic studies demonstrate that age-related vascular problems are often associated with poor cognitive and functional outcomes in older adult populations^1,2^. Age-related differences in brain tissue atrophy have been shown to reflect individual differences in cardiorespiratory fitness^3–5^.

Aging is typically associated with changes in arterial function, such as decreases in the elasticity of cerebral arteries (i.e., arterial stiffening). Arterial stiffening leads to faster pulse wave velocity^6^, which, in turn, modifies the shape of the arterial pulse, due to increased temporal overlap between the forward wave generated by the heart pulsation and the corresponding backward wave generated by peripheral resistance at the level of the muscular arteries and arterioles^7^. This leads to larger and sharper pulse waves in older adults.

It should be noted, however, that although generated within the arteries, the cardiac pulsatile motion can propagate through tissue and, in principle, also be observed in the brain parenchyma. This motion occurs throughout the brain, and while the term “pulsatility” can be used in different ways, here we refer to pulsatility as the degree to which a signal occurs at the frequency band of cardiac pulsation. Cerebral pulsatility has been the subject of increasing interest for its relevance in neurological aging and disease as well as for resting-state fMRI studies. Cerebrovascular aging studies of pulsatility have found that age-related arterial stiffening may produce changes in dampening effects on pulsatile flow into capillaries^8,9^. In addition to providing an understanding of the physiology of vascular wall changes with age, pulsation around blood vessels can have a critical role in glymphatic flow. Cranial CSF flow in the interstitial space surrounding arteries is facilitated by cardiac pulsation, and this process depends on local arterial wall integrity as well as systemic cardiovascular parameters such as blood pressure^10^. This perivascular CSF pumping is the basis for glymphatic clearance, which may impact the ability to remove accumulated amyloid-β associated with Alzheimer disease^11^. For these reasons, major strides have been made in the development of methods to estimate, directly measure, and parameterize pulsatile fluid dynamics in the brain^12^.

Vascular aging has been measured in the brain using diffuse optical methods that characterize cerebral pulsation waveforms using a high temporal resolution and high sensitivity to oxy/deoxy hemoglobin^13,14^. Previous studies have shown that local changes in arterial elasticity are associated with changes in brain volume in the same regions^15^. However, optical imaging approaches have some limitations when it comes to depth of penetration, which limits studies of changes in vascular reactivity in deeper cortices and subcortical structures in cerebrovascular aging. Translation of pulsatile imaging to MRI would enable analysis of deeper cortical regions such as the hippocampus.

Methods such as fMRI offer superior spatial resolution and whole-brain imaging to examine vascular elasticity and reactivity. BOLD has also been used with CO_2_ inhalation or with breath-holding tasks to measure cerebrovascular reactivity, an important measure for vascular health and aging^16–18^. However, fMRI studies with a typical repetition time (TR) of 1 second lack the temporal resolution required to resolve the cardiac pulse waveform to examine passive vascular elasticity. At typical resting-state fMRI sampling rates ranging between 0.5 to 1 Hz, a resting heart rate of 60 beats per minute would exceed the Nyquist limit even for the primary harmonic^19^, and the pulse signal would be aliased at other frequencies. However, characterization of the waveforms can be achieved with the fMRI data after it has been retrospectively cardiac-gated to align with the cardiac phase^20–22^.

Previous studies modified fMRI protocols with shorter TR to achieve better characterization of cardiac pulsations in the brain, motivated by both the methodological benefits of this characterization (more effective removal of physiological noise from resting-state fMRI data) as well as the ability to analyze the pulsations to understand cerebrovascular dynamics. Although physiological noise is often a nuisance signal in resting state fMRI, several studies have targeted the physiological variations in BOLD fMRI data as a signal with useful information^20,23–28^. For example, one study used a 250 ms TR with EPI to cover a small volume (four 5-mm slices) of the brain to measure the average amplitude around the heartbeat frequency in normal appearing white matter (WM) and found that pulsatility differences occur between young adults, old adults, and old adults with small vessel disease^25^. To cover a significant volume at the same TR, simultaneous multislice imaging can be used, which involves simultaneously sampling multiple slices to increase coverage within a TR^20^.

In addition to directly imaging the pulsatile waveforms with fast BOLD fMRI sequences, the pulse plethysmograph signal (PPG) measured simultaneously with the resting state fMRI scan can be used to determine heart rate and cardiac phase for retrospective realignment^20,21,27,29,30^. These methods require merging data over many heartbeats to obtain sufficient samples across the temporal profile of the cardiac waveform. With this approach, previous literature has shown a decrease in BOLD magnitude signal in the arteries with systole and localization of the pulse waveform to the major arteries and CSF^20,21,27,31^. Several studies with a retrospective alignment measured pulsatility by fitting 7-term Fourier series to the retrospectively cardiac phase aligned (using PPG) fMRI data and measuring pulsatility as the fit of the curve^29,30^. Some studies use the amplitude of the retrospectively aligned curve as a measure of pulsatility, which can be done after computing percent signal changes^20^, by normalizing to the standard deviation across time for the voxel^21^, or by normalizing to a baseline signal with low pulsatility^22^. When the cardiac frequency is within Nyquist limits, pulsatility can be defined as the average amplitude within a given interval of the heart rate frequency^25^. Using a 0.02 Hz interval, WM hyperintensities associated with small vessel disease exhibited lower pulsatility than normal WM, however older adults showed higher pulsatility in normal WM than younger adults^25^. Both extremes of pulsatility can indicate vascular dysfunction. A U-shaped association was found between the pulsatility index (measured by ultrasound) and trail-making test, where both low and high pulsatility resulted in increases in time to complete the test^32^.

Other novel MRI-based approaches have been able to achieve high temporal resolution to directly measure pulsatile waveforms. A 3D fast method called magnetic resonance encephalography, MREG, uses spatial localization from large receiver array coils to achieve faster functional imaging^31^. MREG is able to achieve the Nyquist sampling of the heartbeat, but suffers from limited spatial resolution^31^. It has been used to detect heart rate harmonic power and links those differences to Alzheimer’s disease^33^. Harmonics are a relevant feature of aortic blood pressure waveforms and have been shown to be related to the biomechanics and stiffness of vessels^34^.

In this work, we develop a sequence to achieve whole brain, high spatiotemporal resolution imaging to properly resolve cerebral pulse signals anywhere in the brain. To accomplish this, we use a partial separability (PS) MRI acquisition and reconstruction^35–37^ on 3D data with full-brain coverage, 2 mm isotropic resolution, with a TR of 65 ms, acquired at 7T. We used 7T as the contribution of physiological signal relative to thermal noise in the fMRI timeseries increases with field strength^38^. With this short TR, there are 15 timepoint samples over the course of a typical heartbeat cycle at 60 bpm. We demonstrate that the partial separability fMRI (PS-fMRI) sequence allows for pulsatility detection robustly at the first 4 cardiac harmonics, though the amplitude of the oscillations drops substantially at the highest harmonics. With full-brain coverage, we compare pulsatility in WM, gray matter (GM), and cerebrospinal fluid (CSF) and within the reliable pulse regions. To demonstrate the validity of the pulsatility signal and its sensitivity to arterial stiffening, we replicated previous findings of increased pulsatility in older adults relative to younger adults^9,22,25^.

Sex is often included as a variable in examining vascular changes with age due to sex-related differences in neurovascular aging^39^. Pulsatility and pulsatile damping (decrease in pulsatility index as wave travels) has been shown to have sex-related differences with phase-contrast MRI^40^. Therefore, we included sex as a covariate in our analysis and examined relationships between sex and harmonics of pulsatility in our data.

## 2 METHODS

### 2.1 Participants

All procedures in this study were approved by the Institutional Review Board of the University of Illinois at Urbana-Champaign. 28 volunteers were recruited to this study from a pool of participants that had previously taken part in studies in the laboratory. Participants were asked if their health status had changed from the last data collection appointment and were then screened for MRI contraindications. They were instructed to refrain from exercise, caffeine, or use of illicit or vasodilatory drugs for 24 hours prior to their appointment. Upon arrival for their scan, volunteers gave informed consent to participate and were screened for cognitive impairment using the Montreal Cognitive Assessment^41^. Of the 28 participants’ data collected, 2 were not used in the analysis phase due to missing physiological files, 2 were not used due to missing anatomical data, and 2 were unusable due to corrupted PPG, leaving 22 subjects before outlier removal, which removed 1 more participant as explained in Section 2.5.2. Demographics for the subjects prior to outlier removal are shown in Table 1.

**Table 1.** Demographic information for sample before outlier removal.

|  | **All** | **Older** | **Younger** | ***p*** | **Female** | **Male** | ***p*** |
| --- | --- | --- | --- | --- | --- | --- | --- |
| *N* | 22 | 14 | 8 | -- | 10 | 12 | -- |
| **Age *M(SD)*** | 53.0 (17.6) | 65.6 (5.0) | 31.0 (3.0) | < .001 | 50.3 (17.2) | 56.3 (18.2) | .445 |
| **MoCA *M(SD)*** | 28.1 (1.6) | 27.9 (1.5) | 28.6 (1.9) | 0.404 | 27.5 (1.3) | 28.7 (1.8) | .102 |

### 2.2 Scanning procedure

Scans were performed using a Siemens Magnetom Terra 7 T scanner at the Carle Illinois Advanced Imaging Center. Participants were safety screened an additional time by an MRI scan technician prior to scanning. For physiological data collection, participants were outfitted with the MRI scanner provided respiration belt across their chest and pulse oximeter on one finger, interfaced through the Siemens wireless physiological monitoring hardware. The scan session lasted approximately one hour, with the fast fMRI scan lasting 10 minutes 24 s, acquired in the first half of the scan session. Participants were asked to stay awake and stare at a white cross on a black background for the duration of the scan.

### 2.3 PS-fMRI Sequence and Reconstruction

The PS-fMRI sequence leveraged the partial separability framework^35^, where a spatiotemporal image is represented by a sum of the product of a small number of spatial and temporal basis functions. This low-rank acquisition and reconstruction approach allows for the design of a sequence with interleaved temporal navigator signals from a small, repeated region of k-space from which the temporal bases are estimated. This is intermixed with spatial acquisitions that sample all of k-space in a multi-shot approach for the corresponding spatial maps. This enables the estimation of space and time with separate but interleaved acquisitions to create a high spatiotemporal image series^42,43^. For the navigator in the PS-fMRI sequence, a short (1 shot of a 12-shot spiral design, only kz=0, 4.65 ms readout) spiral-in readout was used. This was coupled with a spiral-out imaging acquisition, with a 2-shot per kz-plane with acceleration factor of 2 (4-shots designed, 2 shots 180° rotated) readout with a 24 cm field of view with a matrix size of 120 and 60 kz-planes yielding an isotropic resolution of 2 mm. Only one shot in one kz-plane was acquired for imaging data per acquisition (coupled with a fixed temporal navigator) with sequential sampling of kz-lines. The flip angle was 20° and fat saturation was used with a chemical shift selective module^44^. In the 10-minute, 24-second-long acquisition, 9600 temporal image frames are acquired for the full brain 3D dataset. Image reconstruction proceeds following a similar PS model reconstruction as found in^43^, namely performing singular value decomposition of the temporal navigator signals, choosing the top 35 components, and estimating the associated spatial bases to provide a full framerate reconstruction of the timeseries. In addition, a sensitivity map and field map estimation sequence was acquired just prior to the PS-fMRI sequence to assist in reconstruction.

Due to residual coherency in sampling with a non-random kz line order, some residual temporal spikes were present in the timeseries, distributed across the brain volume. To eliminate this spiking, the fBIRN small QC phantom (EZfMRI, Chicago, IL) was scanned with the same protocol demonstrating the same sharp noise spikes. Ignoring the first and last 250 points in the 9600-point spectrum, the signal was thresholded, resulting in a mask of 160 frequency locations (out of 9600) that are removed from the spectrum of the temporal basis in each participant’s data to eliminate the sharp frequency spikes, see Supplementary Figure S1 for visual inspection of the filter’s performance.

### 2.4 Data processing

#### 2.4.1 Physiological data

Physiological data output from the scanner was cleaned to remove header information and non-measurement codings, then aligned to the beginning of the fMRI scan for analysis.

#### 2.4.2 Anatomical data

Anatomical T1-weighted images for functional registration were collected using a Magnetization Prepared Two Rapid Acquisition Gradient Echo (MP2RAGE) sequence^45^ with scan parameters TR = 4.53 s, TE = 2.26 ms, inversion times = 750/2950 ms, flip angles = 4° and 5°, voxel size = 0.7×0.7×0.8 mm^3^. MP2RAGE images were denoised using the LayNii package^46^ and brain extraction was performed using FSL BET^47^. Anatomical data were segmented by tissue type using FreeSurfer 5.3.0^48^ to obtain GM, WM, and CSF masks. Time-of-flight angiography was also employed to obtain maps of vasculature (TR = 8.3 ms, TE = 2.46 ms, flip angle = 20°, voxel size = 0.5×0.5×0.5 mm^3^).

#### 2.4.3 Functional data

Registration of fMRI to MNI space was performed with advanced normalization tools in *python*^49^. The sensitivity map reference images were used as the PS-fMRI reference as they were taken during the same scan and had the same dimensions as the fMRI scan. Data were linearly registered to anatomical, and SyN transformed (affine then warp) to MNI space. The transformations were saved so they could be applied to results in the native fMRI space of each subject.

### 2.5 Analysis

#### 2.5.1 Pulse reliability

For each subject, the first 300 repetitions were removed to reach steady state. The PS-fMRI data were then broken up into 18 windows of 32.5 s. Each window was analyzed for pulse reliability, a voxel-wise measure of the quality of the pulsatile fMRI signal, after band-pass filtering from 0.1 to 5.5 Hz. Pulse reliability was measured by retrospective cardiac gating the time series into 30 bins according to the position of the time point in the cardiac phase, measured linearly from 0 to 1 for preceding peak in the PPG signal to the next peak, for even and odd heartbeats separately, then determining the coefficient of determination (R^2^) between them, which is similar to the method described by Hermes et al. 2023^20^. Pulse reliability measures the quality of the pulse signal within each voxel and each window. An example of pulse reliability measurements for five voxels are shown for one subject and for one window in Figure 1. The pulse-realigned fMRI signal curves show that voxels corresponding to CSF have a signal with a peak near the PPG systole (cardiac phase 0 and 1)^20^, with a CSF voxel shown in Figure 1b (voxel 1). Large arteries have a dip at systole^20^, which occurs close to the PPG systole, shown in Figure 1c and 1d (voxels 2 and 3). Figure 1e is another example of a voxel likely to reflect arterial signal and Figure 1f shows a voxel with low pulse reliability.

**Figure 1.**
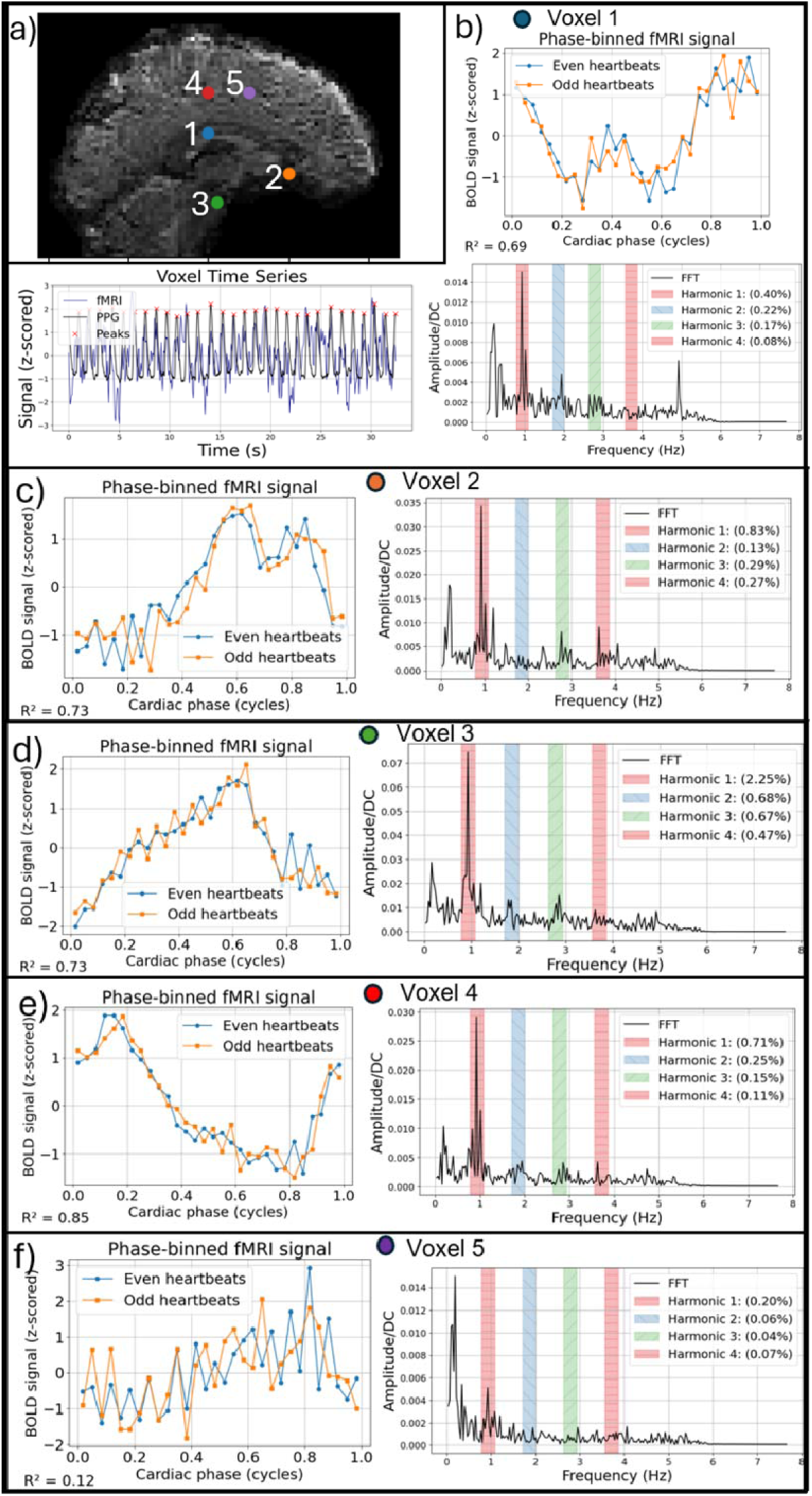
Individual voxels from one subject for one 32.5 second time window. a) Mean fMRI image with labeled voxels; b) Voxel 1 and PPG time series (bottom left), realigned pulse with reliability (top right), and frequency spectra (bottom right); c-f) realigned pulse for voxels 2-5 (left) and frequency spectra for voxels 2-5 (right). Voxels 1-4 have high pulse reliability, indicating a quality heartbeat signal. Voxel 5 has low pulse reliability.

The pulse reliability maps were averaged across temporal windows for each subject by first converting the R^2^ to a z-score using the Fisher z-transform and then averaging across the windows, followed by using the inverse Fisher z-transform to return to R^2^ values for each voxel for each subject. Transformations described in 2.4.3 were applied to each R^2^ map to convert them to MNI space and averaged across subjects, again using the Fisher z-transform, to get a combined average R^2^ in the MNI space. The group reliability map was thresholded at 0.4 to make a pulse reliability mask for restriction of further analysis to voxels with high pulse reliability.

#### 2.5.2 Pulsatility

Pulsatility in the MRI data was measured in a manner similar to Makedonov et al. 2013^25^. Each voxel’s unfiltered FFT magnitude was computed and normalized to the DC component, and the average magnitude within 0.15 Hz of the heart rate, as found from the PPG of the window using *findpeaks* in SciPy, was calculated as the pulsatility for that window. Examples of pulsatility of the harmonics in individual voxels are shown in Figure 1.

The WM, GM and CSF partial volume masks were transformed into the native fMRI space for each subject to compute the percent cardiac pulsatility within each region. The group pulse reliability mask was transformed from MNI to each subject’s native fMRI space. Only voxels within the MNI brain mask and pulse reliability mask registered to each subject’s native fMRI space were included in the analysis. Percent cardiac pulsatility was first averaged across windows for each subject, then averaged within each ROI according to each tissue type mask (using a weighted average by partial fraction). An ordinary least square test was used to assess differences in pulsatility between younger and older adults, using sex as a covariate and treating age group as a categorical variable. To correct for multiple comparisons within each harmonic metric, the false discovery rate method^50^ was applied using multiple tests from *statsmodels.stats.multitest* in *python* with the “fdr_bh” method at alpha = 0.05^51^.

Outlier removal was applied within each subject across temporal windows. Outlier time windows, defined as having percent cardiac pulsatility within any region (GM, WM, CSF) and any harmonic below the first quartile minus 3 times the IQR or above the third quartile plus 3 times the IQR were removed before averaging. Across all subjects, a total of 15 time-windows were removed. The greatest number of windows removed from one subject was 4. Subject outlier removal removed 1 young male subject, leaving 21 total subjects for statistical analysis of the harmonics. The outlier was determined by removing any subject with mean pulsatility across non-outlier windows above the third quartile plus ten times the interquartile range or below the first quartile minus ten times the interquartile range.

To determine whether pulse reliability and pulsatility were localized to vessels, Spearman correlation coefficients were computed between the pulse reliability map and vessels from a previously published atlas^52^. The vessel map was resampled to the 1mm isotropic MNI template and clipped from 0 to 100 (probability 0 to 1) prior to computing Spearman correlations. After applying an MNI brain mask, a pulse reliability mask of R^2^ > 0.05, and vessel (large artery) map mask of probability > 0.05, the rank order Spearman correlation between the pulse reliability map and the vessel probability mask^52^ was computed: 1) within a WM mask where voxels were included if they had a partial fraction in the WM > 0.5 (WM mask) and 2) within a GM mask, where voxels where in included if they had partial fraction in the GM > 0.5. In addition, we computed the correlation between the percent cardiac pulsatility at each harmonic with the pulse reliability within the MNI brain mask. We also calculated the correlation between the percent cardiac pulsatility at the first harmonic and the vessel probability map in masks consisting of WM and GM, including voxels with WM > 0.5 or GM > 0.5.

#### 2.5.3 Correlations with pulse

PPG recordings were smoothed and resampled to the temporal resolution of the PS-fMRI scan and aligned before taking correlations. MATLAB *corrcoef* was used to calculate the Pearson correlation coefficient between the time series of each voxel with the PPG. FMRI time series were linearly detrended via first order polynomial regression prior to computing correlations for each voxel.

## 3 RESULTS

### 3.1 Pulse Reliability

The pulse reliability across all subjects is shown in Figure 2a, showing regions of high-quality pulsatile signal. The vessel map derived from the atlas from Mouches & Forkert 2019^52^ is shown for comparison in Figure 2b. The Spearman correlation coefficients were 0.39 (p < 0.0001) for the WM mask and 0.28 (p < 0.0001) for the GM mask, showing that the regions with reliable pulsatility were generally localized to the vessels. Graphs of the Spearman correlations between pulse reliability and vessel probability for each voxel are shown in Supplementary Figure S2.

**Figure 2.**
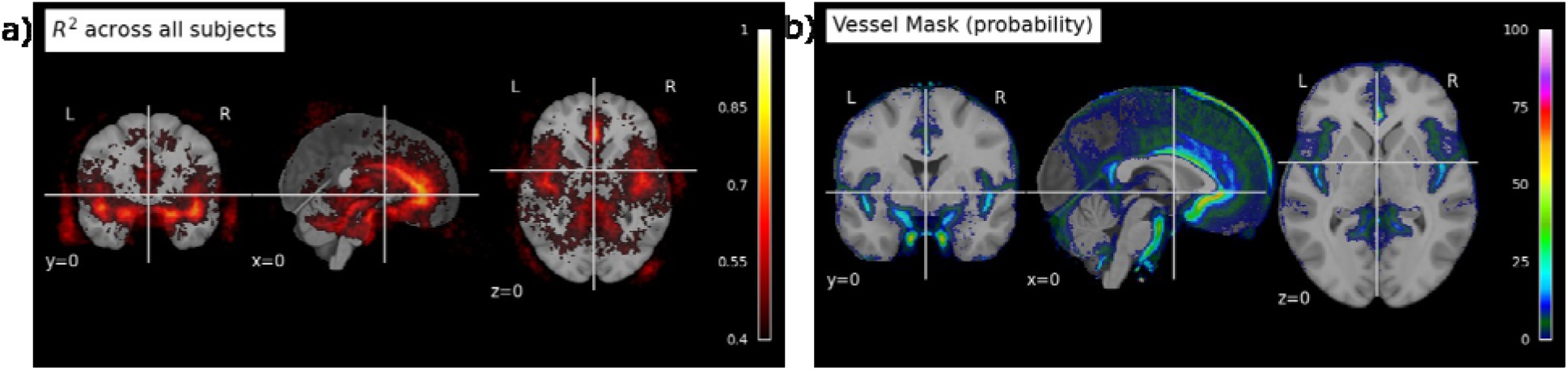
a) Pulse reliability maps across subjects showing R^2^ > 0.4; b) visualization of the vessel map atlas from Mouches & Forkert 2019 ^52^, downsampled to 1mm isotropic to match the MNI template, then clipped from 0 to 100 to get values corresponding to probability.

### 3.2 Pulsatility

A statistical analysis was performed between younger (N =7, aged 25 to 40 years of age) and older (N = 14, aged 55 to 75 years of age) adults. The results for CSF, GM, and WM masks, harmonic, and harmonic ratio *(% cardiac pulsatility of harmonic / % cardiac pulsatility of first harmonic)*, with sex used as a covariate, were calculated for the first 4 harmonics and first 3 harmonic ratios. Significance (*p* < 0.05 after multiple comparisons correction) was found for GM and WM for the first and second harmonics.

The older group had a higher magnitude for all significant differences. Plots for each comparison are shown in Figure 3a for age differences and Figure 3b for sex differences. Males had a larger magnitude in the third harmonic normalized to the first in CSF, GM, and WM. Cohen’s *d*, along with the uncorrected and corrected *p*-value for each comparison are shown in Tables 2 and 3 for age and sex differences, respectively.

**Figure 3.**
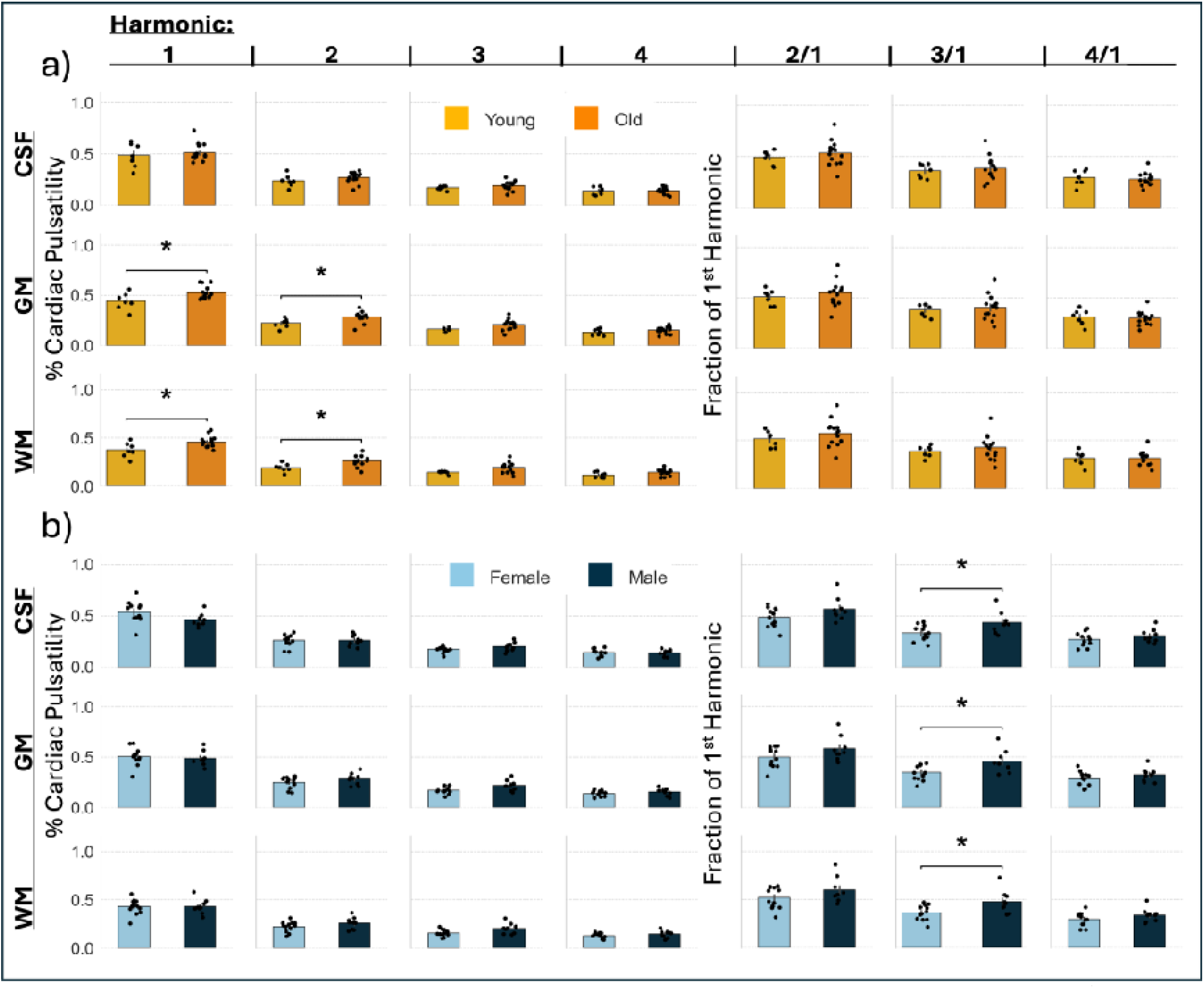
Bar plots representing average % cardiac pulsatility. Group differences are shown for a) young vs old and b) male vs female for 4 harmonics and 3 harmonic ratios. Bars indicate standard errors. Statistical results from OLS with p < 0.05 (FDR correction within each column, separately for male/female and young/old) being labeled significant.

**Table 2.** Age differences in cardiac pulsatility between young (*n =* 7) and old (*n =* 14) adults (FDR per metric across ROIs). The q-value is the p-value after correcting for multiple comparisons. Hn = Harmonic *n*. * indicates significance at _α_ = 0.05.

| | Young<br>$M \pm SE$ | Old<br>$M \pm SE$ | Cohen's $d$ | $p$-value | $q$-value<br>(FDR) |
| --- | --- | --- | --- | --- | --- |
| **CSF** |  |  |  |  |  |
| **Harmonic 1** | $0.487 \pm 0.044$ | $0.514 \pm 0.022$ | 0.28 | 0.297 | 0.297 |
| **Harmonic 2** | $0.237 \pm 0.022$ | $0.269 \pm 0.014$ | 0.59 | 0.225 | 0.225 |
| **Harmonic 3** | $0.169 \pm 0.007$ | $0.191 \pm 0.012$ | 0.57 | 0.353 | 0.353 |
| **Harmonic 4** | $0.139 \pm 0.014$ | $0.141 \pm 0.008$ | 0.06 | 0.885 | 0.885 |
| **Ratio H2/H1** | $0.492 \pm 0.026$ | $0.532 \pm 0.033$ | 0.37 | 0.631 | 0.631 |
| **Ratio H3/H1** | $0.360 \pm 0.025$ | $0.380 \pm 0.031$ | 0.2 | 0.97 | 0.97 |
| **Ratio H4/H1** | $0.293 \pm 0.029$ | $0.277 \pm 0.017$ | -0.24 | 0.434 | 0.757 |
| **GM** |  |  |  |  |  |
| **Harmonic 1** | $0.441 \pm 0.032$ | $0.527 \pm 0.017$ | 1.21 | 0.011* | 0.022* |
| **Harmonic 2** | $0.222 \pm 0.017$ | $0.286 \pm 0.014$ | 1.27 | 0.023* | 0.035* |
| **Harmonic 3** | $0.162 \pm 0.006$ | $0.208 \pm 0.014$ | 1.06 | 0.060 | 0.090 |
| **Harmonic 4** | $0.133 \pm 0.012$ | $0.154 \pm 0.009$ | 0.66 | 0.246 | 0.369 |
| **Ratio H2/H1** | $0.507 \pm 0.029$ | $0.550 \pm 0.034$ | 0.37 | 0.630 | 0.631 |
| **Ratio H3/H1** | $0.377 \pm 0.024$ | $0.403 \pm 0.033$ | 0.24 | 0.958 | 0.970 |
| **Ratio H4/H1** | $0.308 \pm 0.029$ | $0.295 \pm 0.018$ | -0.18 | 0.504 | 0.757 |
| **WM** |  |  |  |  |  |
| **Harmonic 1** | $0.375 \pm 0.029$ | $0.457 \pm 0.016$ | 1.25 | 0.015* | 0.022* |
| **Harmonic 2** | $0.192 \pm 0.016$ | $0.260 \pm 0.014$ | 1.33 | 0.018* | 0.035* |
| **Harmonic 3** | $0.142 \pm 0.006$ | $0.193 \pm 0.014$ | 1.13 | 0.043* | 0.090 |
| **Harmonic 4** | $0.116 \pm 0.011$ | $0.144 \pm 0.009$ | 0.86 | 0.127 | 0.369 |
| **Ratio H2/H1** | $0.516 \pm 0.031$ | $0.575 \pm 0.037$ | 0.48 | 0.448 | 0.631 |
| **Ratio H3/H1** | $0.388 \pm 0.022$ | $0.427 \pm 0.034$ | 0.36 | 0.730 | 0.970 |
| **Ratio H4/H1** | $0.316 \pm 0.029$ | $0.316 \pm 0.020$ | -0.0 | 0.787 | 0.787 |

**Table 3.**
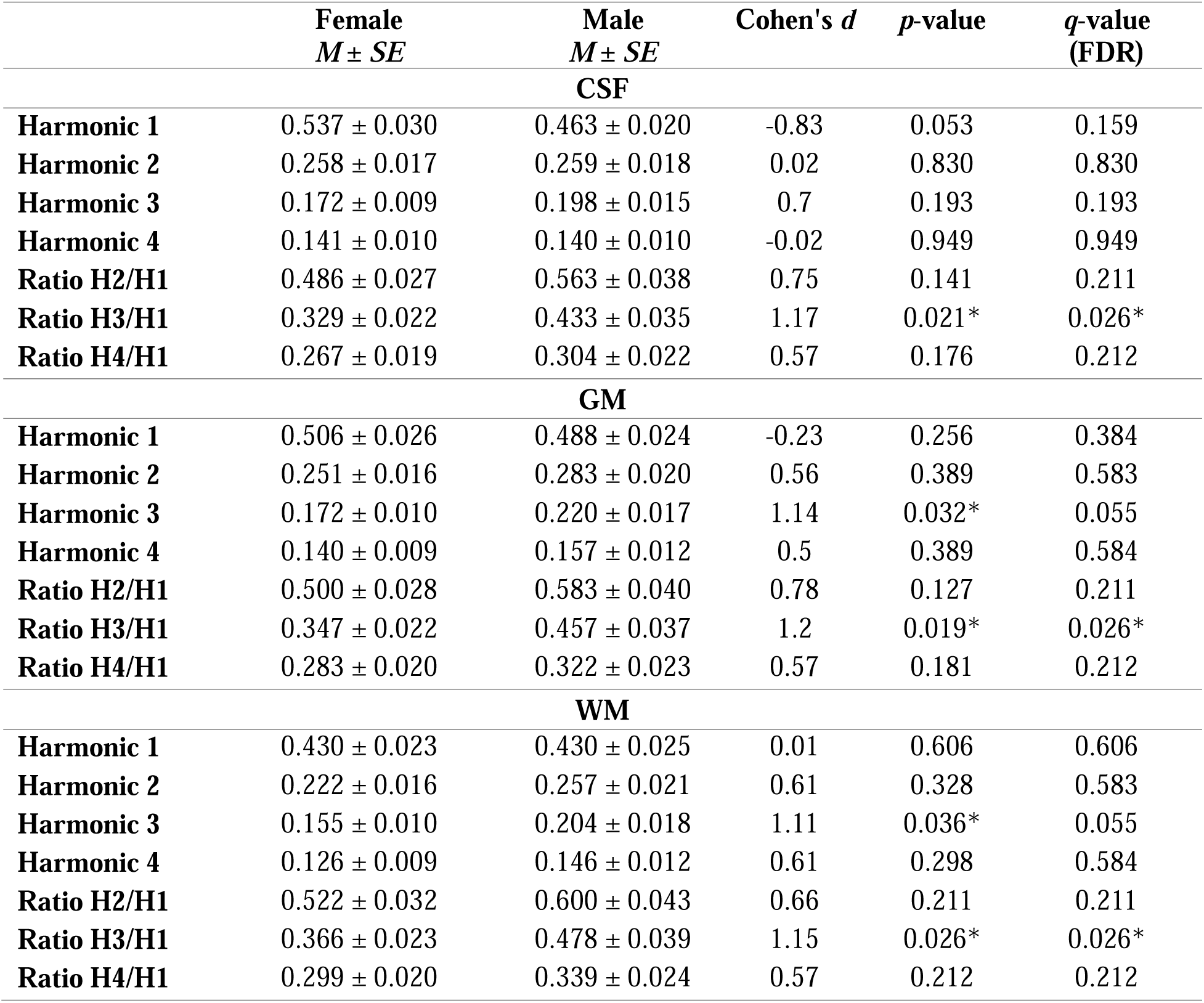
Sex differences in cardiac pulsatility between female (*n =* 12) and male (*n =* 9) participants (FDR per metric across ROIs). The q-value is the p-value after correcting for multiple comparisons. Hn = Harmonic *n*. * indicates significance at _α_ = 0.05.

The average pulsatility map averaged across all 21 subjects after transforming each subject to MNI is shown in Figure 4 for all harmonics. Percent cardiac pulsatility within reliable (R^2^ > 0.4) regions in the brain was correlated to the pulse reliability as shown by significant rank order Spearman correlations for all harmonics (correlations equal to 0.22, 0.18, 0.13, 0.10 for harmonics 1, 2, 3, and 4, respectively with *p* < 0.0001 for all harmonics). The first harmonic was correlated to the vessel probability in a mask consisting of voxels with partial volume measures either WM > 0.5 or GM > 0.5 with Spearman correlation of 0.32 (p < 0.0001). Plots of the pulse reliability and vessel probability vs percent cardiac pulsatility for each voxel used in the correlation are shown in Supplementary Figure S3.

**Figure 4.**
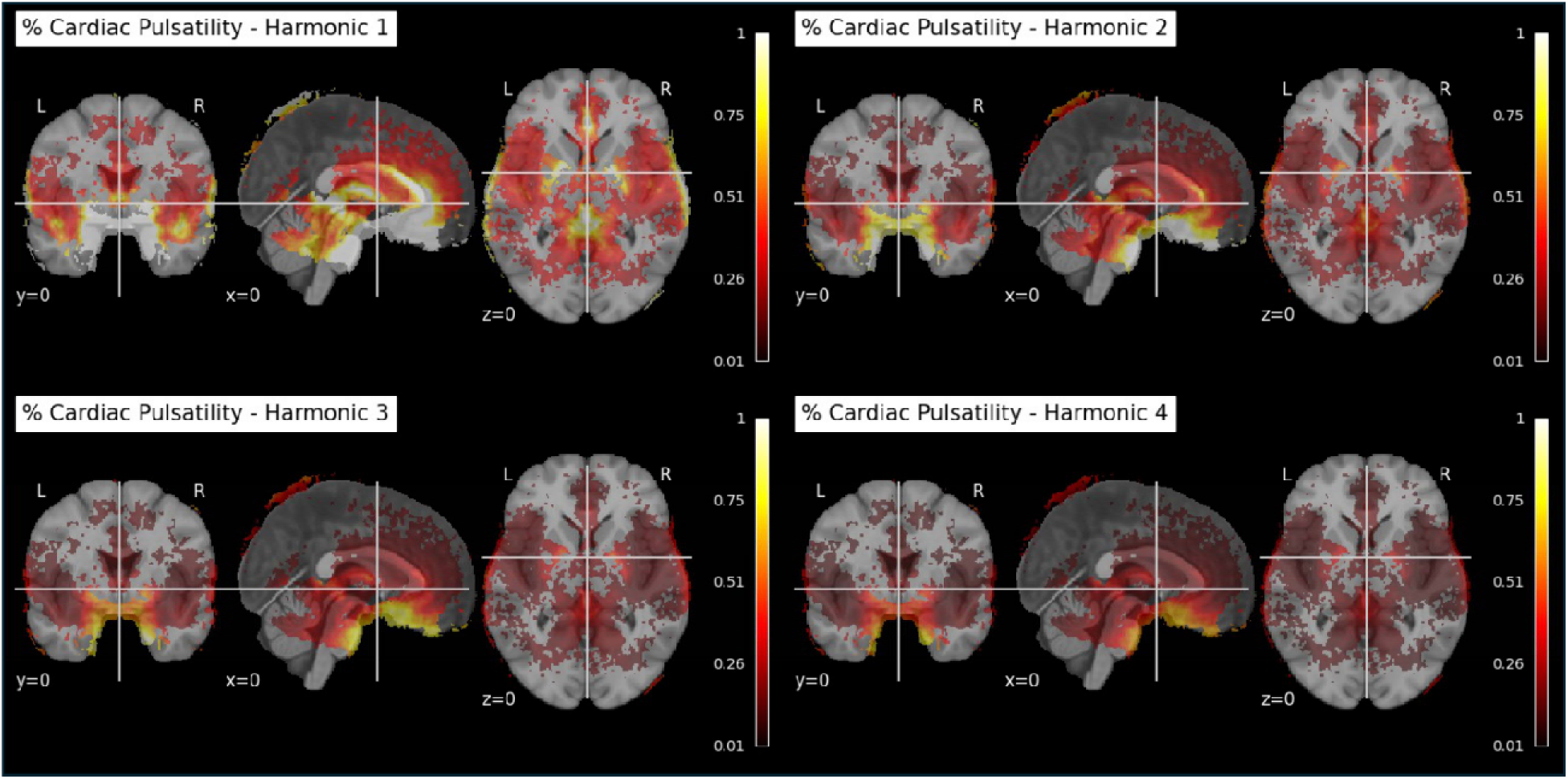
Percent cardiac pulsatility maps for 1^st^, 2^nd^, 3^rd^, and 4^th^ harmonics, clockwise from top left respectively. MNI brain mask and pulse reliability mask R^2^ > 0.4 applied.

### 3.3 Correlation Maps

Maps of correlation of the magnitude of pulsations in each voxel with the PPG-measured pulse are shown in Figure 5. The correlation maps follow the typical vascular structure for major arteries, with higher correlations concentrated near the circle of Willis, bilateral middle cerebral arteries, and anterior cerebral artery, showing high localization of signal to vascular areas.

**Figure 5.**
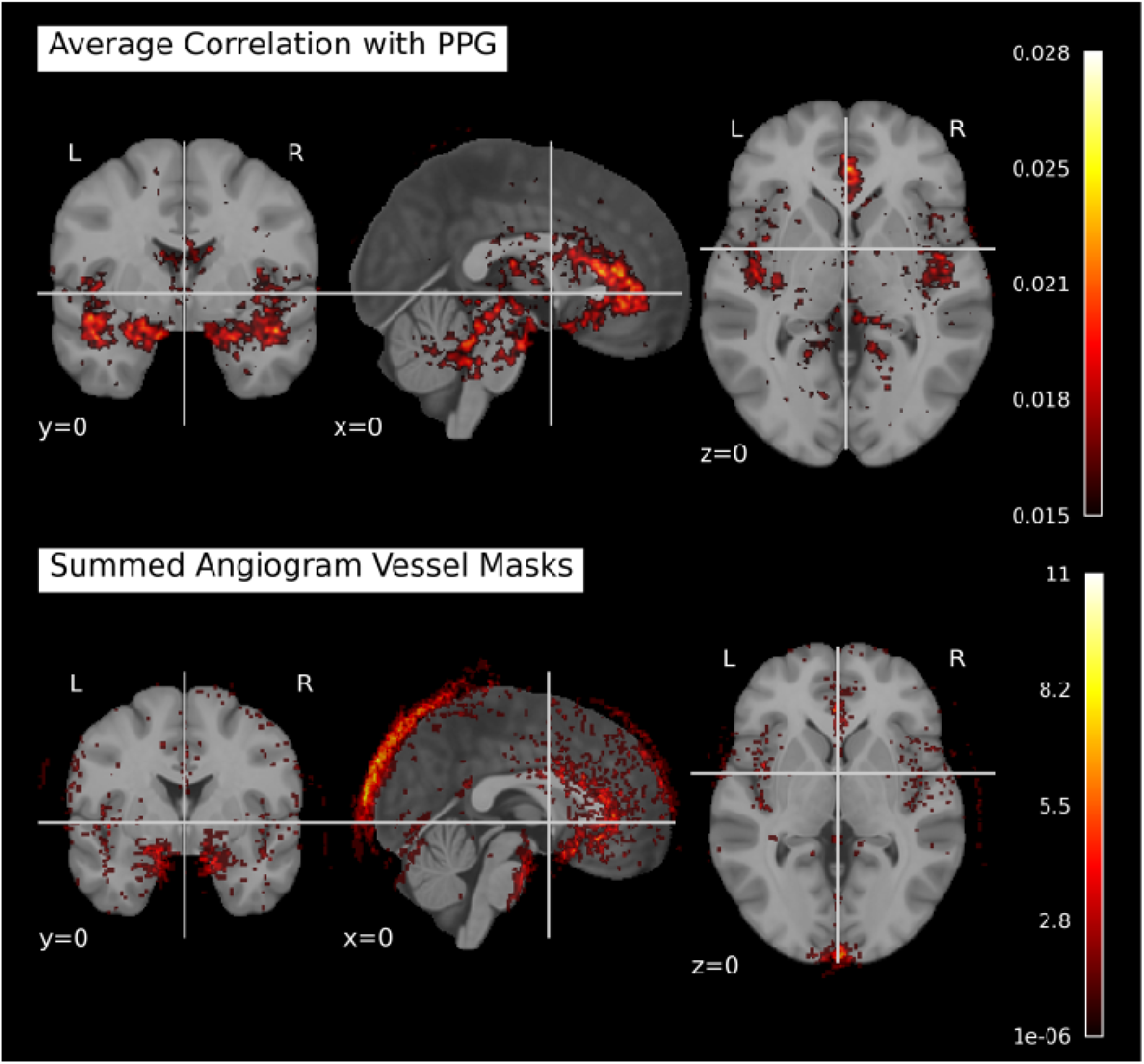
Upper panel shows magnitude Pearson correlations for each voxel (normalized) with PPG in MNI space averaged across all subjects thresholded for correlations > .015. Lower panel shows sum of all angiograms masked to vasculature (>200) transformed to functional and then to MNI space. Taken together these images show the overlap between the areas with relatively high degree of correlation with pulse and cerebroarterial vasculature as captured by angiography. The highest correlations are primarily clustered near the base of the brain in the area of the circle of Willis (top right axial), middle cerebral artery (middle, coronal), and anterior cerebral artery (lower left, sagittal). Reliability of this pattern of correlations is demonstrated in Supplementary Figure 4.

## 4 DISCUSSION AND CONCLUSION

### 4.1 PS-fMRI can reliably measure pulsation throughout the brain

The PS-fMRI sequence achieved 2-mm isotropic, whole brain coverage with a TR of 65 ms to enable visualization of pulsation waveforms throughout the brain. Using this TR, we were able to resolve the primary cardiac pulsation signal and 4 of its harmonics. These pulsations are reliable and are localized around the vascular structures in the brain. The odd-even heart beats provide significantly correlated waveforms and percentage pulse amplitude is correlated with pulse reliability in high-reliability regions. This indicates that the PS-fMRI sequence is suitable for further work resolving temporal features of the pulsatile signal throughout the brain.

### 4.1 Age- and sex-related pulsatility

Our results agree with previous literature^9,22,25^ by showing an age-related increase in pulsatility, measured as the DC-normalized average magnitude around (0.3 Hz interval) the heart-rate frequency throughout the brain. Previous literature found increased pulsatility, measured within 0.02 Hz around the heartbeat frequency, in older adults in normal appearing WM within a limited acquisition of four 5 mm thick slices^25^. Another study defined pulsatility as *(maximum of pulse amplitude – minimum of pulse amplitude) / median of a baseline signal* within the circle of Willis (single-slice) and found increased pulsatility within older adults^22^. This is consistent with previous PC-MRI studies showing a higher pulsatility index, defined as *(systolic blood flow rate - diastolic blood flow rate) / (mean blood flow rate)*, in older adults^9^. This work furthers these previous findings by achieving total brain coverage, fast acquisition, and high spatial resolution. The PS model can be used to expand the ROIs accessible when measuring pulsatility with short TR.

In addition, there are significant differences in the CSF, GM and WM masks, where males show a larger third harmonic ratio normalized to the first (p < 0.05). In the literature, a similar observation was made using tonometry to measure the pulse waveform at the wrist^53^, with males having greater third harmonic ratio spectral power^54^. This result may reflect pressure and blood volume measures, similar to PS-fMRI measures. The displacement of the vessels is thought to contribute to the BOLD fMRI signal^29^, which could explain the appearance of this sex difference with PS-fMRI.

The localization of the pulse reliability signal and pulsatility to the vessels, as indicated by the significant correlations, is consistent with previous results^20,27^. The pulse reliability correlations with all 4 harmonics shows every harmonic considered contributes to pulse reliability. The pulse reliability in the sagittal sinus is low, while other studies at lower field strength and higher TR show pulse reliability within the sagittal sinus^20^. This may be due to the decrease in T2* at 7 T from the high deoxyhemoglobin in veins.

The dicrotic notch, which is affected by age, may be related to the ratio of the second harmonic normalized to the first, and has been shown to change with age in PPG waveforms^55^. In the PS-fMRI data, there was not a clear dicrotic notch after the retrospective realignment, and Figure 3 shows that the ratio of the second harmonic normalized to the first failed to reach significance between age groups. The realignment assumed a linear increase in phase from one peak of the PPG to the next. A method that uses a nonlinear phase increase has been shown to enable detection of a dicrotic notch in fMRI signals^21,22^. This new method may yield more detailed cardiac pulsatility curves that allow detection of localized differences in the dicrotic notch. Modified realignment could expose features related to vascular stiffness, as is done with PPG^56,57^ and fNIRs^14^ features.

### 4.2 Limitations

There are several limitations to this study. While the use of a short TR provided the temporal resolution to record over each pulse cycle, shorter TRs also reduce SNR due to the shortened time for longitudinal magnetization recovery^23,58–60^ and increase the size of the data. The large amount of data (9600 time points of 120×120×60 voxels per subject) takes a longer time to process and requires a workstation with large memory, as each fMRI data file itself is > 60 GB.

There are also several limitations with the current sequence. First, whole brain coverage at 65 ms temporal resolution with dynamic imaging requires imaging a spatiotemporal model that leverages redundancy in the image series. With a limited model order, this captures most of the signal in the image series but may lose subtle cardiovascular pulsations that are hard to separate from noise. Further, due to the sampling scheme of serial ordering of kz-encodings, some spike noise was coherent in our sampling. Although we do not expect cardiac pulsations to be time-locked to our acquisition, the removal of a small number of spike components (only 160 out of 9600 spectral locations) could result in differential removal of information depending on the specific subject’s heart rate.

Retrospective cardiac gating applied in this study requires an adequate PPG recording. 2 subjects were dropped due to poor PPG data. Methods that don’t require a reference cardiac sensor can use data from subjects with poor recordings^61^, and would increase statistical power if an accurate alignment can be recovered.

### 4.3 Future directions

Diffuse optical imaging pulse studies of the brain have been shown to reveal information about arterial stiffness, an important descriptor of cerebrovascular health related to aging and fitness^14,62,63^. Future work in ultra-fast MRI includes exploring the heterogeneity of the pulse waveforms observed throughout the brain, the physiological basis of this heterogeneity, and what we can infer about brain health using such indicators. It is important to note that there are distinctions in the pulse waveform recorded from optical methods and flow-based fMRI, and successful interpretation of these waveforms depends on understanding the basis for signal differences between modalities^64^.

The PS-fMRI approach could also be used to test how TR affects either the pulsatile signal or the neural signal of interest through subsampling methods, where preliminary results show differences^37^. A similar analysis has been done with MREG^65^. Finding differences in physiological noise correction from TR and age groups could offer new insight into potential fMRI confounds.

### 4.4 Conclusion

PS-fMRI is a new fMRI imaging method that uses the PS model to sample whole brain BOLD-based fMRI, with 2 mm isotropic resolution at an ultra-fast TR of 65 ms. The pulsatile signal can be reliably detected across 4 harmonics of the heart cycle, is localized to the CSF and vessels, and can detect significant age and sex differences. This could enable the study of age-related changes to the elasticity of blood vessels throughout the brain.

## ACKNOWLEDGMENTS

A seed grant from the Beckman Institute Biomedical Imaging Center supported the 7 T data collection. This work was also partially supported by NIA grants R01AG059878 and RF1AG062666 to M. Fabiani and G. Gratton, and by a Carle-Beckman Fellowship to Dema Abdelkarim. This work was conducted in part at the Biomedical Imaging Center of the Beckman Institute for Advanced Science and Technology at the University of Illinois Urbana-Champaign (UIUC-BI-BIC).

## Data Availability Statement

Data and code for the current study are available from the corresponding author (BS) upon reasonable request.

## TABLE AND FIGURE CAPTIONS

Supplementary Figure S1. a) Phantom frequency spectrum and filter. After averaging the voxels within the phantom ROI, the magnitude of the FFT of the time series was taken. b) The filter from masking out spikes above the 1e-5 (AU) threshold. c) The first temporal basis function FFT magnitude for one subject and d) after filtering by multiplication with the filter in b.

Supplementary Figure S2. Hexagonal bins showing density of voxels. a) Vessel probability vs pulse reliability within R^2^ > 0.05 mask, and vessel probability > 0.05 mask for within the WM > 0.5 mask and b) within the GM > 0.5 mask, showing the density of voxels in hexagonal bins. _p_ is the Spearman correlation coefficient and _p_ is the p-value.

Supplementary Figure S3. Hexagonal bins showing density of voxels. a-d) Pulse reliability vs average magnitude at the harmonic for all 4 harmonics within the MNI brain mask. e) Spearman correlation (_p_) and p-value (_p_) between mean % pulsatility and vessel probability within a mask that is (GM > 0.5 or WM > 0.5).

