## Supplemental Information for "Ultrafast fMRI detects age-related changes in harmonics of cardiac pulsations in the brain at 7 T"

**Supplementary Information**

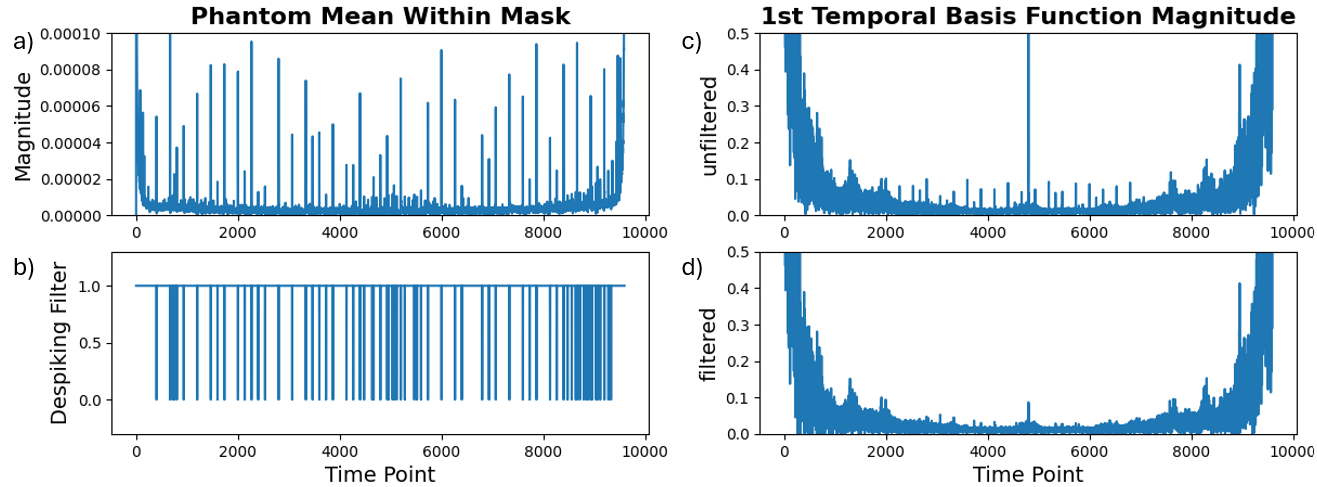
 Supplementary Figure S1. a) Phantom frequency spectrum and filter. After averaging the voxels within the phantom ROI, the magnitude of the FFT of the time series was taken. b) The filter from masking out spikes above the 1e-5 (AU) threshold. c) The first temporal basis function FFT magnitude for one subject and d) after filtering by multiplication with the filter in b.

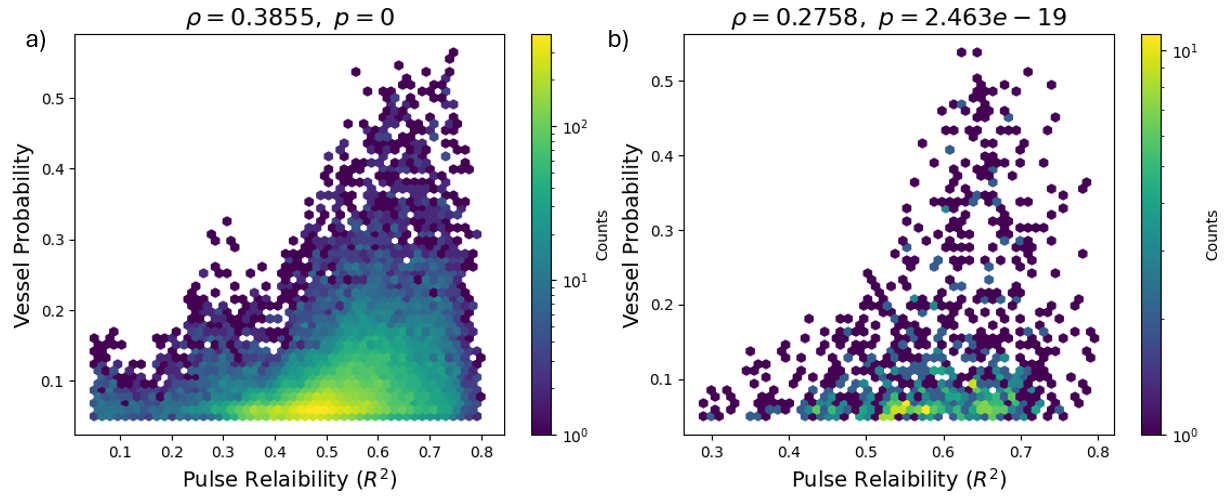
 Supplementary Figure S2. Hexagonal bins showing density of voxels. a) Vessel probability vs pulse reliability within R^2^ > 0.05 mask, and vessel probability > 0.05 mask for within the WM > 0.5 mask and b) within the GM > 0.5 mask, showing the density of voxels in hexagonal bins. 𝜌 is the Spearman correlation coefficient and 𝑝 is the p-value.

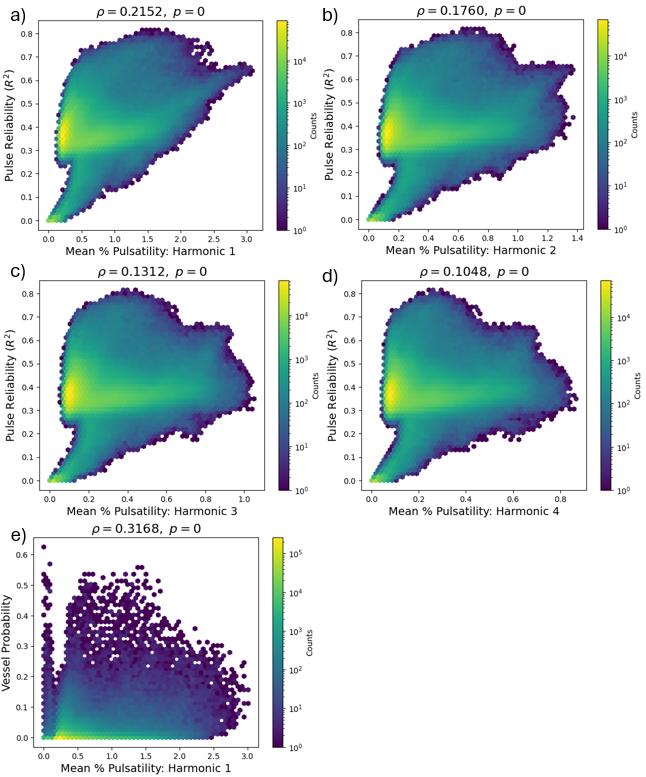

Supplementary Figure S3. Hexagonal bins showing density of voxels. a-d) Pulse reliability vs average magnitude at the harmonic for all 4 harmonics within the MNI brain mask. e) Spearman correlation (𝜌) and p-value (𝑝) between mean % pulsatility and vessel probability within a mask that is (GM > 0.5 or WM > 0.5).
